# Normal human lymph node T follicular helper cells and germinal center B cells accessed via fine needle aspirations

**DOI:** 10.1101/757534

**Authors:** Colin Havenar-Daughton, Isabel G. Newton, Somaye Y Zare, Samantha M. Reiss, Min Ji Suh, Farnaz Hasteh, Shane Crotty

## Abstract

Germinal centers (GC) are critically important for the maturation of the antibody response and the generation of memory B cells, which are the basis for long-term protection from pathogens. Germinal centers only occur in lymphoid tissue, such as lymph nodes, and are not present in blood. Therefore, cells of the germinal center, including GC B cells and GC T follicular helper (T_FH_) cells, are not well-studied in humans under normal healthy conditions, due to the limited availability of healthy lymph node samples. We used a minimally invasive, routine clinical procedure, lymph node fine needle aspirations (LN FNAs), to obtain lymph node cells from healthy human subjects to establish benchmarks of GC cells under noninflammatory conditions. This study of 50 lymph nodes demonstrates that human LN FNAs are a safe and feasible technique for immunological research, and defines benchmarks for human GC biology. The findings indicate that assessment of the GC response via LN FNAs will have application to the study of human vaccination, allergy, and autoimmune disease.

## INTRODUCTION

The efficacy of the vast majority of human vaccines depends on antibody (Ab) responses. Our knowledge of the immunological processes controlling the generation of neutralizing Abs is insufficient, and therefore progress on how to optimize neutralizing antibody generation after immunization has been largely empirical (1, 2). Induction of protective Abs by vaccination depends on complex interactions of the immune system. CD4 T cell and B cell responses are essential for developing vaccine-elicited neutralizing Abs but remain poorly characterized for any vaccine.

Within lymph nodes (LNs), germinal centers (GCs) are sites of intricate B cell and CD4 T follicular helper (T_FH_) cell interaction (2, 3). GC responses are necessary for the development of Ab affinity maturation and memory B cells. GC B cells and GC-T_FH_ cells are not present in blood and, therefore, can only be studied with lymphoid tissue samples (4). Whole LN surgical excisions are rarely performed in humans for research purposes. When they are done, it is most commonly in the context of a disease state (5–7). Tonsils have served as a widespread alternative source of human GC cells, but the fact that tonsils have exposed surfaces to the environment leave unknowns about phenotypes of GC cells in more sterile LN environments. Additionally, the inability to directly quantify GC activity (GC B cells and GC-T_FH_ cells) after specific immunizations or other interventions has remained a vexing ‘‘black box’’ problem for many human immunology studies, as most studies depend entirely on blood samples.

Fine needle aspirations (FNA) are an alternative means of sampling LNs. FNAs were first described in the medical literature nearly 100 years ago. It is a simple outpatient procedure with minimal risk that is generally non-disruptive to the LN architecture. As such, FNAs are widely used in clinical medicine for the diagnostic assessment of abnormal LNs and masses (8–10). They have been used sporadically in immunology and infectious disease research (1, 11–13). To assess whether LN FNAs would be a useful immunological research technique for the study of GC B cells and GC-T_FH_ cells, we conducted two extensive studies in non-human primates (14, 15). We first determined that LN FNAs provide a representative sample of the lymphocyte subsets in the LN (14) and that LN FNAs reproducibly provide sufficient cells for immunological analysis. We then used LN FNAs to explore GC responses in primates after protein immunization. In over 1000 procedures, LN FNAs were used to quantitatively measure GC activity in the LNs and provided valuable correlates of Ab responses. In these studies, HIV neutralizing Abs were generated by candidate HIV vaccine immunogens at rates much higher than previous studies. The generation of these HIV neutralizing Abs strongly and specifically correlated with GC activity uncovered by LN FNA sampling. Most importantly, the immunological knowledge gained from the LN FNAs guided changes in vaccination schedules and strategies, which resulted in a much higher success of the vaccine (15, 16). Based on the strength of these findings, we have now assessed the feasibility of implementing LN FNAs as a research tool in humans for the study of LN-specific cells and processes, such as GC-T_FH_ cells and GC B cells. We have found that ultrasound-guided FNAs can successfully and safely recover cells from human axillary and inguinal LNs, including GC-T_FH_ and GC B cells.

## RESULTS

### Successful and safe ultrasound-guided LN FNAs in healthy human donors

LN FNAs in healthy humans presented a number of challenges that need to be addressed before it can be widely used as a research tool. Are putatively quiescent LNs identifiable and technically feasible to sample by FNA? Does LN FNA cause undue discomfort in subjects? Are sufficient cells recovered for analysis? Does blood contamination during LN FNAs confound analysis? Are T cells, B cells, GC-T_FH_ cells, and GC B cells detectible?

Successful recovery of cellular material from FNAs performed on enlarged LNs for clinical diagnosis is routine. However, FNAs performed on putatively quiescent LNs from healthy human subjects are far more challenging. We recruited a cohort of 23 healthy human subjects for LN FNAs (**Table 1**). The cohort contained 18 female and 5 male subjects. The average age was 28.9 years old (range 19-58), with an average BMI of 24.5 (range 18.3-32.9).

**Table 1:** Human subject characterist cs

|  |  | Total (n = 23) | Mean | SD | Range |
| --- | --- | --- | --- | --- | --- |
| Gender | Female | 18 (78%) |  |  |  |
|  | Male | 5 (22%) |  |  |  |
| Age (years) |  |  | 28.9 | 10.1 | 19-58 |
| BMI |  |  | 24.5 | 3.5 | 18.3-32.9 |

Superficial LNs are present in the human axillary and inguinal regions and are not easily palpable in most healthy human subjects. In accordance with standard clinical practice, we used ultrasound (US) to identify and target axillary and inguinal LNs and to avoid adjacent vessels and other structures (**Figure 1A**). LNs generally measured 1 cm in the greatest long axis dimension (**Figure 1B**). Normal LNs demonstrate a fatty echogenic hilum and a hypoechoic cortex, which usually measures less than 2-3 mm. On average, the visual identification of LNs in healthy human donors was judged as being moderately challenging by the physician performing the procedure (**Figure 1C**). Discrimination of normal LNs on US improved with experience, per operator feedback. LN FNA samples were attempted at 2-4 regional LN sites (bilateral inguinal and bilateral axillary sites) per donor for a total of n=50 LN FNA attempts.

**Figure 1.**
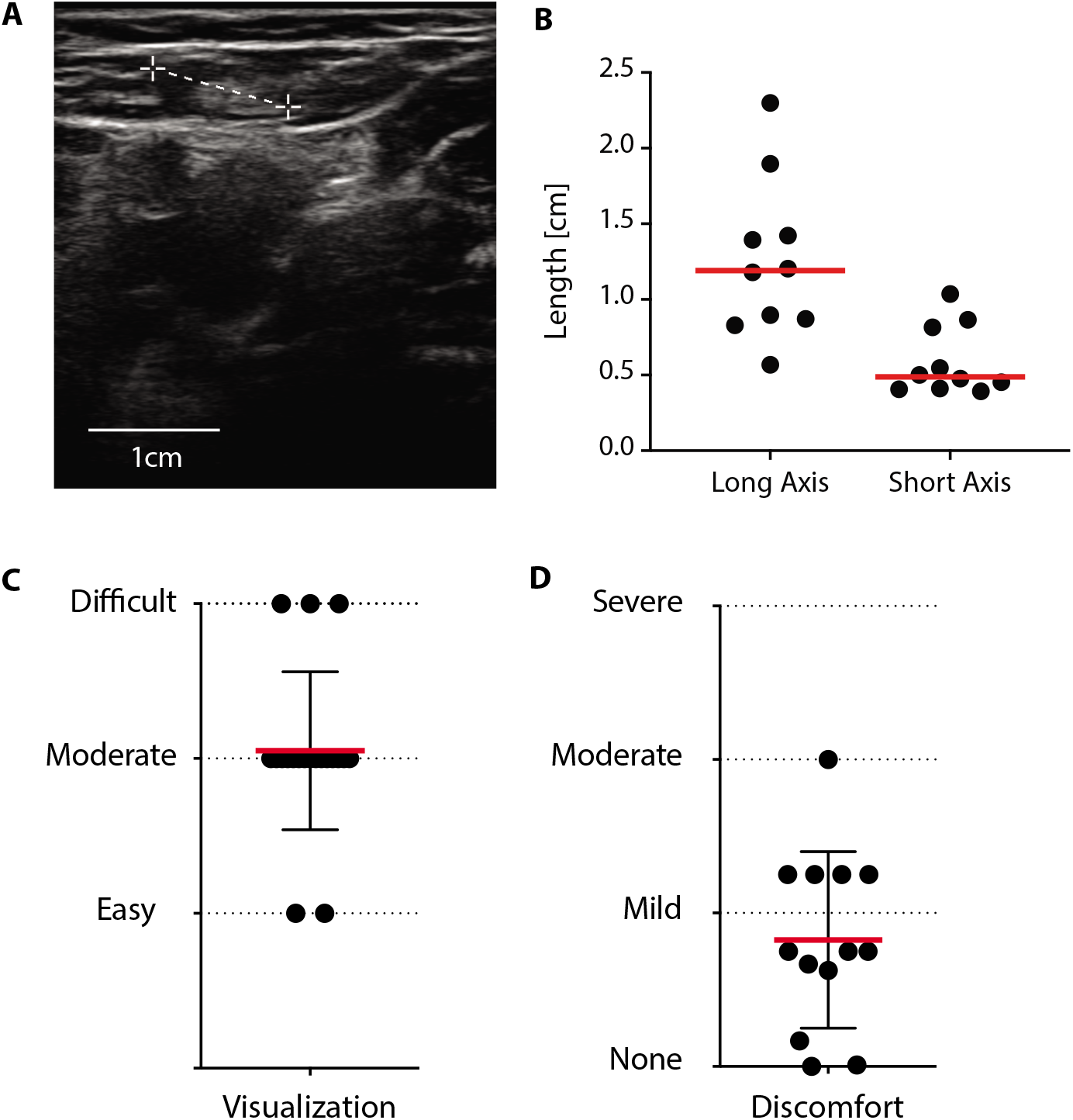
Ultrasound guided LN FNAs in healthy human donors is well-tolerated. A) Ultrasound image of LN. B) Graph of LN sizes, measured via ultrasound. C) Visualization score of LNs images by ultrasound. D) Graph of donor reported discomfort during the LN aspiration procedure. Red bars indicate geometric means.

Healthy subjects tolerated LN FNAs well. Because obtaining LN FNAs from healthy human subjects could be limited by discomfort, we asked each subject to rate their degree of discomfort for each FNA performed at each LN site. Most subjects reported no discomfort to mild discomfort (**Figure 1D**). There were no major complications observed in the study. Overall, US-guided FNA represents a safe, feasible, well-tolerated method for sampling LNs in healthy subjects for research purposes..

### Cell recovery of human LN FNAs in healthy human donors

The number of cells recovered by LN FNAs is a key criterion for its utility as a diagnostic and research tool. Technological advances in highly multiparameter flow cytometry and CyTOF analysis, flow cytometric cell sorting, and single cell transcriptional analysis have reduced the number of cells need to obtain biologically meaningful data. Including all LN FNA attempts, the median cell recovery for axillary LN FNA samples was 219,000 cells; the median cell recovery for inguinal LN FNA samples was 513,000 cells (**Figure 2**). The number of cells recovered ranged between 4,000 and 7,900,000, a level of variability that could be expected from a technically challenging sampling procedure. Six-eight percent (34 of 50) of LN FNA samples contained >100,000 cells. In summary, the majority of LN FNA samples from heathy human subjects provided sufficient cell numbers for flow cytometry analysis.

**Figure 2.**
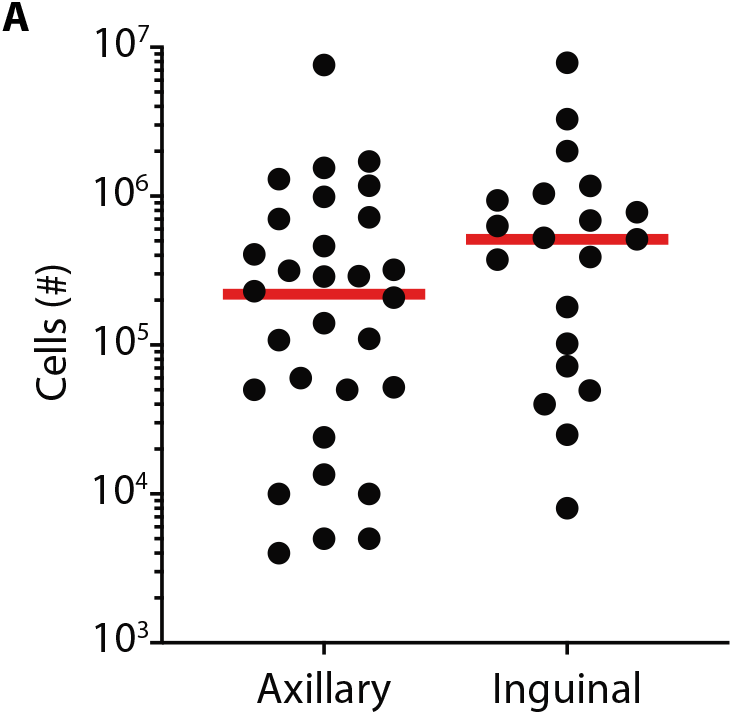
Human LN FNA cell recovery. A) Cell recovery of lymphocytes from human LN FNAs. Red bars indicate geometric means.

### LN FNA sample purity

We used flow cytometry to assess the cellular composition of the LN FNA samples. We considered how to best assess the degree of blood contamination in the LN FNA samples. Granulocytes are the most common leukocyte in whole blood (40-70%)(17), but rare in LN tissue; therefore, we assessed granulocyte frequency as a measure of blood contamination in the LN samples. The LN FNA samples contained a mean frequency of 4.3% granulocytes (range 0.3 to 40%, **Figure 3A-B**), indicating low levels of blood cell contamination. Next, we gated on the lymphocyte population (**Figure 3A**) and expression of the CD45 marker and a live-dead stain (**Figure 3C**) to discriminate live hematopoietic cells from other cells or debris. The median frequency of live CD45+ cells from LN FNA samples was >95% (**Figure 3D**), indicative of healthy samples.

**Figure 3.**
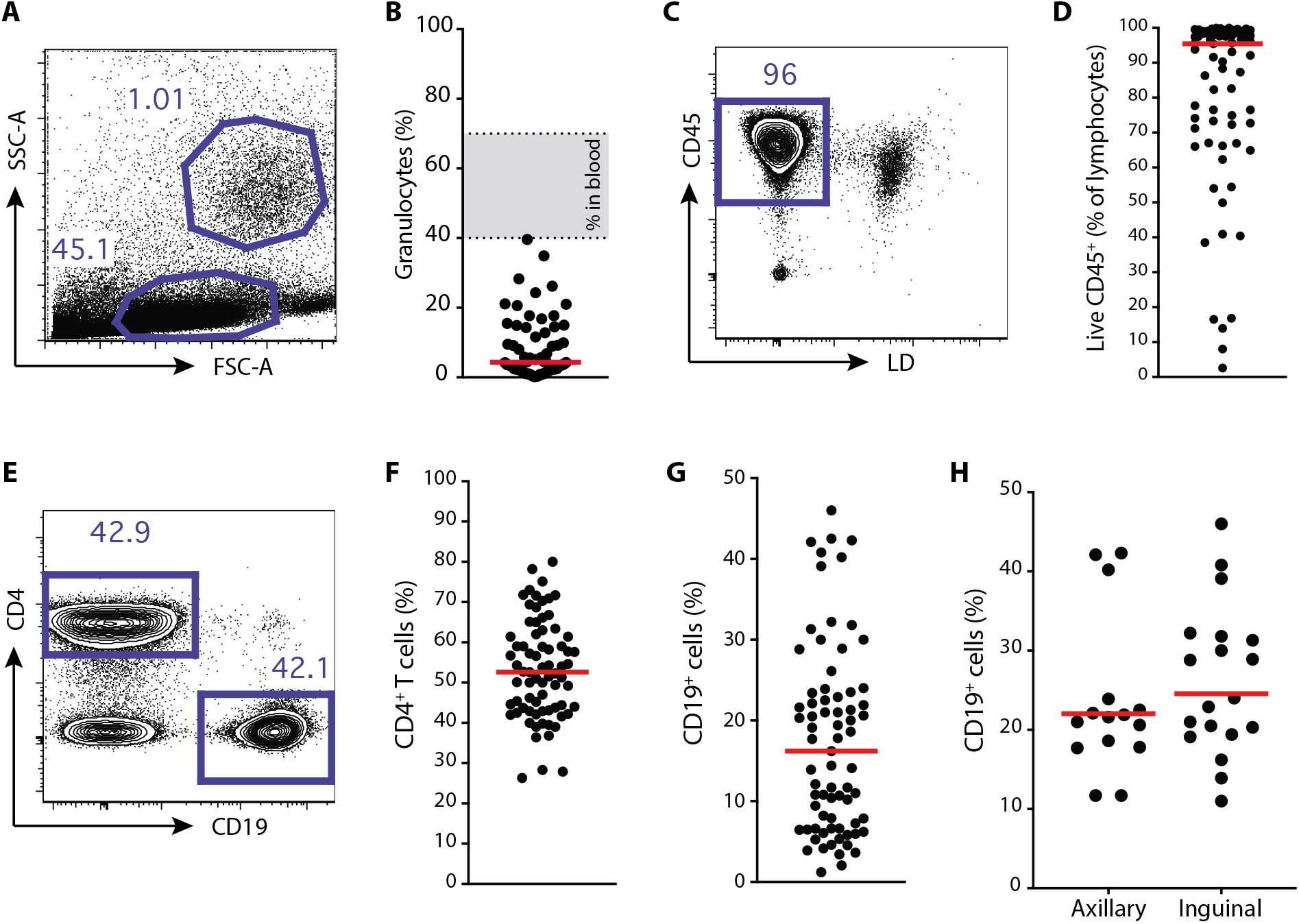
Live T and B cells recovered from healthy human LN FNA samples with minimal blood cells. A) Representative flow plot of lymphocyte and granulocyte populations. B) Graph of granulocyte percentage of total acquired events. Gray shaded area represents the frequency range of granulocytes found in blood from normal healthy subjects (ref17). C) Representative flow plot of live, CD45^+^ lymphocytes. D) Quantification of live CD45^+^ lymphocytes. E) Representative flow plot of CD4^+^ T cell and B cell gating. F) Quantification of CD4^+^ T cell percentages of live lymphocytes. All samples G) Quantification of CD19^+^ B cell percentages of live lymphocytes. All samples. H) Quantification of CD19^+^ B cell percentages of live lymphocytes. Successful samples. Red bars indicate geometric means.

Among live CD45^+^ lymphocytes, populations of CD4 T cells and B cells were gated (**Figure 3E**). The mean frequency of CD4 T cells in human LNs was 52.3% (**Figure 3F**), which was similar in both axillary and inguinal LNs (**Figure S1**). We noted that the frequency of CD19^+^ B cells did not have a normal distribution and had a lower than expected mean value, with many below 10% (**Figure 3G**). Separately, the data by LN location revealed differences in the frequency of CD19^+^ B cell recovered (**Figure S1**). We hypothesized the differences were due to technical, rather than biological, variation, as axillary LN FNAs are smaller and generally more challenging to visualize and sample. We next developed criteria (see Methods) to gauge the success of the LN FNAs, which included consideration of the total cell recovery of live cells and frequencies of lymphocytes, granulocytes, and B cells. Using these criteria, we found similar frequencies of CD19^+^ B cells in inguinal LNs (mean of 24.6%) and axillary LNs (mean of 22.1%) of high quality samples (**Figure 3H**). Furthermore, we were able to identify populations of CD8 T cells, NK cells, and plasmacytoid and conventional dendritic cells within LN FNA samples (**Figure S2**). Overall, live T and B cells were recovered from healthy human LN FNA samples with minimal blood contamination, with similar frequencies in inguinal and axillary LNs.

### FNAs technique impacts LN sample quality

LN FNA sample quality is a key factor for subsequent immunological analysis. Therefore, we evaluated the data based on subject age, subject BMI, and the physician performing the LN FNA. We found that subject age and BMI did not dramatically affect cell recovery of successful LN FNA samples (**Figure 4A-B**). Furthermore, subject age and BMI did not correlate with LN FNA success, within the ranges tested (**Figure S3**). One physician had greater success and higher cell recovery when performing inguinal LN FNAs as compared to axillary LN FNAs (**Figure 4C**). A second physician had substantial success with axillary LN FNAs (**Figure 4C**). We also tested if the use of aspiration during the FNA increased cell yield. We found that aspiration consistently increased the total lymphocyte recovery by LN FNA (3-fold, p=0.026, **Figure 4D**). Cell subset recovery was not affected by the use of aspiration during the LN FNA (**Figure S3**). In summary, the performing physician and application of aspiration during the LN FNA, but not subject age or BMI, affected sample success and cell recovery in heathy human subjects.

**Figure 4.**
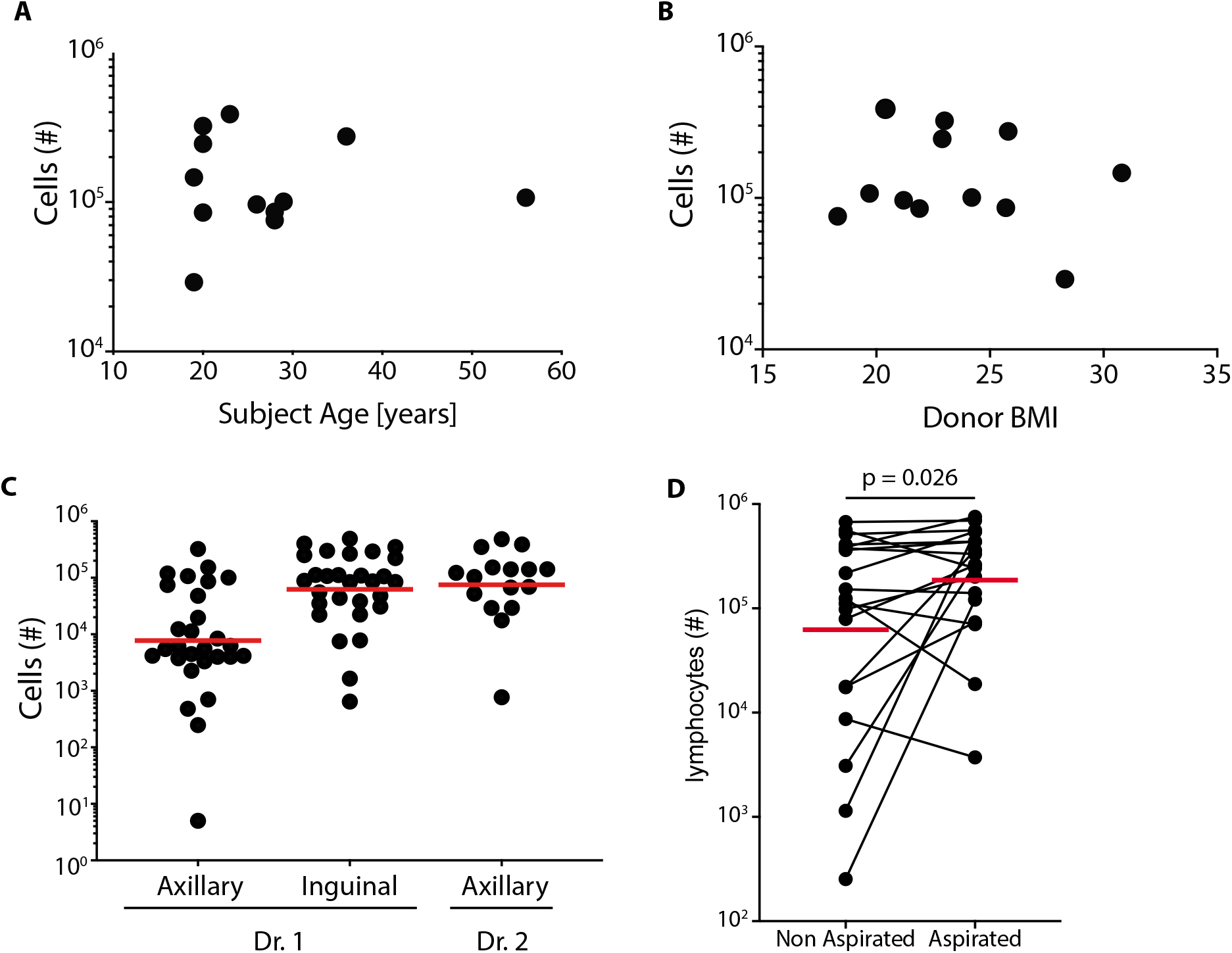
Variables affecting LN FNA sampling. A) Cell yield of live lymphocytes versus donor age. Successful samples. B) Cell yield of live lymphocytes versus donor BMI. Successful samples. C) Cell yield as a function of sampling site and physician. D) Cell yield as a function of with or without aspiration. Two-tailed Wilcoxon matched-pairs signed rank test. Red bars indicate geometric means.

### Quantification of human LN GC-T_FH_ cells and GC B cells by LN FNA

Insights into human Tfh cells and antibody responses have been gleaned from studies of peripheral blood, but direct assessment of normal GC cells in human LNs has been rare. Here, we identified and quantified human LN GC-T_FH_ cells among CD4 T cells (Figure 5A) and GC B cells among CD19^+^ B cells (**Figure 5B**). GC-T_FH_ cells express high levels of CXCR5 and PD-1 (14, 18), and GC B cells were identified using CD20 and CD38. GC-T_FH_ cells and GC B cells are activated cells indicative of ongoing antigen-specific immune responses. Therefore, superficial LNs from healthy human subjects would be expected to have low frequencies of GC cells. GC-T_FH_ cell frequencies in donor LNs ranged from 0.1% to 2.6% (mean 0.4%, **Figure 5C**). GC B cell frequencies in donor LNs ranged from 0.2% to 6% (mean 0.7%, **Figure 5D**). These frequencies were substantially lower than generally observed in tonsils, which can contain 36% GC-T_FH_ of CD4^+^ T cells and 38% GC B cells of total B cells (18). GC-T_FH_ cells and GC B cells from successful LN samples were similar between physicians performing the LN FNAs (**Figure 5F-G**). Interestingly, one subject reported to have received an immunization one month prior to the LN FNA procedure; the LN FNA revealed one of the highest frequencies for GC B cells (3.1%) and GC-T_FH_ cells (1.4%) in the cohort, suggesting that human vaccine responses may be measurable in draining LNs by FNA (Figure 4C and 4D, green point). In conclusion, GC-T_FH_ cells and GC B cells were detectable in LN FNAs at low frequencies in superficial LNs from healthy human subjects.

**Figure 5.**
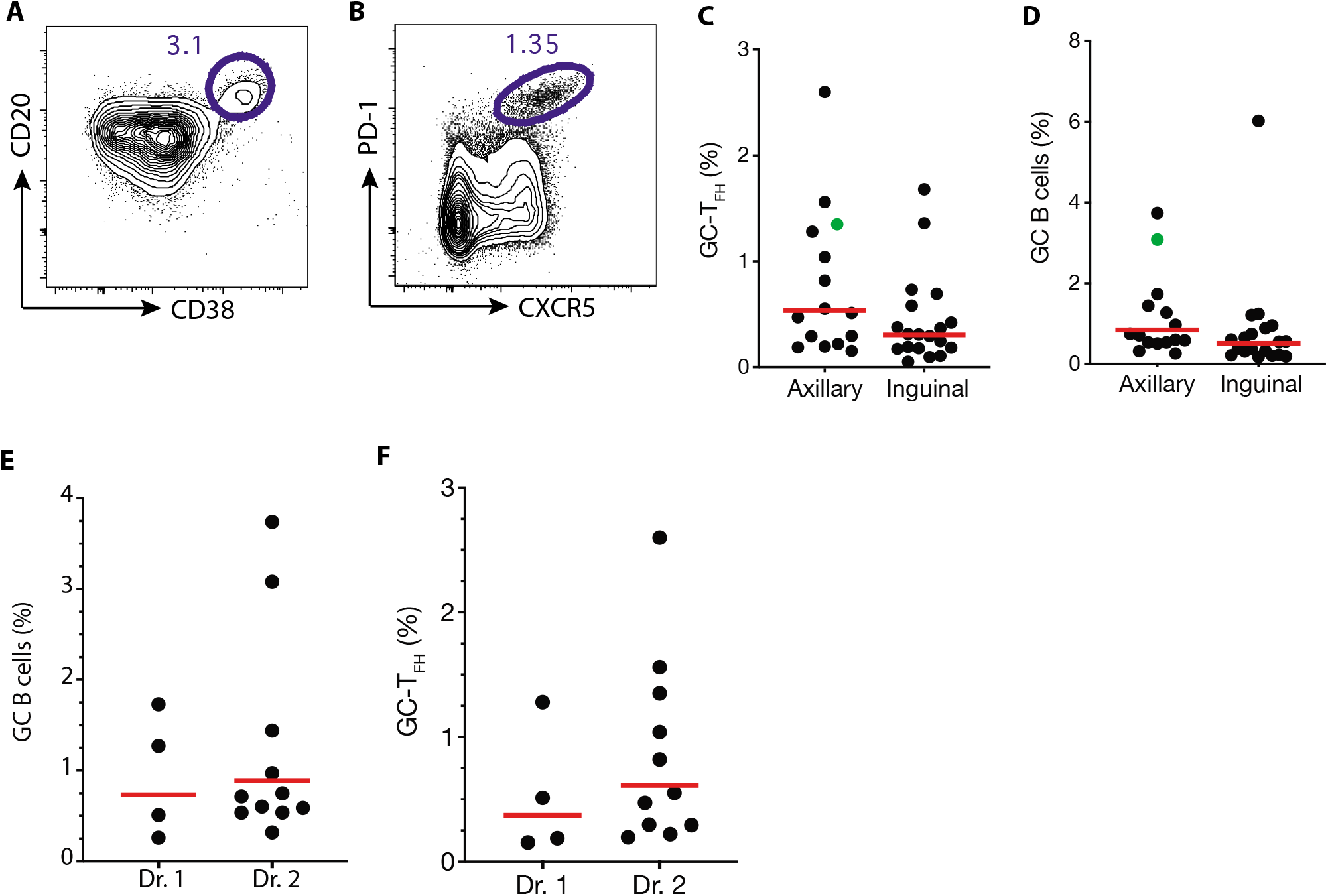
Quantification of human LN GC-T_FH_ cells and GC B cells by LN FNA. A) Representative flow plot of GC B cell gating post immunization in a draining LN. Gated on CD19^+^ B cells. B) Representative flow plot of GC-T_FH_ cell gating post immunization in a draining LN. Gated on CD4^+^ T cells. C) Quantification of GC-T_FH_ % of successful LN samples. Green point, subject reported to have received an immunization one month prior to the LN FNA procedure. D) Quantification of GC-T_FH_ % of successful LN samples. Green point, subject reported to have received an immunization one month prior to the LN FNA procedure. E) Quantification of GC B cell frequency of successful attempts as a function of physician. F) Quantification of GC-T_FH_ cell frequency of successful attempts as a function of physician. Red bars indicate geometric means.

## DISCUSSION

US-guided LN FNAs provide unparalleled access to cells of the LN and allow for the study of GC biology. Because healthy LNs are typically small (1cm in long axis dimension) and quiescent, sampling them by FNA required practice, even for expert operators. To define the quality control criteria for a successful sample we considered numerous factors. Granulocyte frequency was considered a sign of blood contamination due the high frequency of granulocytes found in peripheral blood. Total B cells were expected to be at higher frequencies in LN than in blood. LN-specific cell types such as GC-T_FH_ cells and GC B cells were indicative of successful LN sampling, although these cell types are rare in noninflammatory conditions. Total cell yield also must be considered as a success criteria, as low cell recovery can skew results. Considered together, successful sampling of the LN via FNA should provide a sample of at least 50,000 total cells with at least 5,000 total B cells, a low granulocyte frequency (<10%), and the presence of defined GC-T_FH_ cells and GC B cell populations, even if rare.

US-guided FNAs represent a safe and feasible method for sampling LNs in healthy subjects for research purposes. US guidance is recommended to identify these LNs, which are not usually palpable. US also guides sampling directly from the cortex, where the GCs reside, and helps to reduce complications associated with potential puncture of adjacent structures. On ultrasound, normal LNs demonstrate a fatty echogenic hilum, sometimes with a feeding vessel that is detectable on color Doppler imaging. The hypoechoic cortex of normal lymph nodes usually measures less than 2-3 mm. Reactive and pathologic lymph nodes demonstrate thicker cortices. Because it can be challenging to discriminate lymph nodes with very thin cortices from the surrounding fat and muscle, bidimensional US assessment is recommended.

LN samples are procured from the cortex, with care to avoid the hilum and the vessels it contains. Adequate LN samples may be procured by FNAs using a small needle (27 G) under real-time gray scale US guidance with a high frequency linear probe. A longitudinal approach of the needle with respect to the transducer allows full visualization of the needle during each pass to increase yield and avoid adjacent structures and blood vessels. Once inside the cortex, the needle is rapidly moved back-and-forth in order to accumulate cells within the needle by capillary action. Aspiration may be applied to increase the cell count. While this variation in technique can be associated with greater blood contamination and been reported to degrade the quality of the sample though no significant differences in sample quality were observed in the current study between aspiration and non-aspiration samples from the same LN. Major complications after FNA are exceptionally rare, especially when guided by US. No major complications were observed in the current study.

This study of 50 lymph nodes demonstrates that human LN FNA is a safe and feasible technique for immunological research, and defines benchmarks for human GC biology. The finding indicate that assessment of the GC response via LN FNAs will have application to the study of human vaccination, allergy, and autoimmune disease. Multiple human vaccine clinical studies are currently incorporating LN FNAs into their study design for clinical trials (19), representing rapid adoption of this innovative immunological approach.

## METHODS

### Human subjects

LN FNAs and blood draw protocols were approved by the UCSD Institutional Review Board and accepted by the La Jolla institute Institutional Review Board under a reliance agreement covering the approved protocols. All human subjects gave informed consent before entering the study. Individuals were compensated for their time and effort in the study.

Study inclusion criteria included subjects over the age of 18 years, males or non-pregnant, non-nursing females, self-reported to be feeling well and healthy on day of donation, weighing at least 85 pounds for blood draw, BMI less than 30, a hematocrit count between 38-54%, or hemoglobin level of at least 12.5 g/dl, resting pulse within range of 50-100 bpm, systolic blood pressure of 90-180 mmHg and a diastolic blood pressure of 50-100 mmHg, willingness to provide consent to obtain and store genetic data, and ability to provide signed informed consent. Study exclusion criteria included history of anemia, presence of significant cardiovascular disease, systemic diseases including, but not limited to, uncontrolled diabetes, renal disease, liver disease, malignancy, infection, or coagulopathy, thrombocytopenia, inability to provide informed consent, a hematocrit of <38% or >54%, or hemoglobin level >12.5 g/dl, a pulse outside of the 50-100 bpm range, a systolic blood pressure <90 or >180 mmHg and/or diastolic blood pressure <50 or >100mmHg, blood thinners or aspirin within the last 5 days, age less than 18 years, pregnancy, lactation. In addition, we recommend excluding subjects with breast implants or rotator cuff discomfort from LN FNAs at the axillary site.

Major complications after FNA are exceptionally rare, especially when guided by US. The most common minor complication is access site bleeding, which can be easily controlled with compression, particularly the absence of a coagulopathy or blood thinners. Rarely does a patient develop a hematoma. Infection at the biopsy site is a rare complication when sterile techniques are employed. Also rare is the inadvertent puncture of a tiny nerve radical that is undetectable on ultrasound, resulting in radiating pain or numbness. No major complications were observed in the current study.

### Ultrasound (US)-guided LN FNA procedure

One of two qualified physicians performed the ultrasound (US)-guided procedure with the aid of a nurse or qualified clinical staff. Standard of care was used during all FNAs, including sterile techniques and equipment. Superficial LNs at the axillary site or inguinal site were targeted. Axillary and inguinal LNs were sampled initially but only axillary LNs were biopsied for the second half of the study given future plans to compare lymph nodes post immunization, and the deltoid is the most common site for vaccination. First, the overlying skin was cleaned with iodine or other antiseptic solution and the subject covered with a sterile drape. The ultrasound probe was covered with a sterile sleeve. Real-time US (Philips HD 15 Revision 3.0.1) was used during the procedure to locate LNs and to guide the FNA. Local anesthesia was administered with 1% lidocaine. A maximum of 4 LNs were sampled, with each LN sampled up to 4 times (passes). Each pass was performed with a new, sterile needle, which was moved back-and-forth rapidly within the cortex of the lymph node. Needle size (22 gauge, 25 gauge, or 27 gauge) and syringe size (3 or 5 ml) were dependent on LN FNA operator. In the cases in which aspiration was applied, slight negative pressure was generated by withdrawing the plunger of the syringe during sampling. LN FNA samples were expelled into 0.5 ml of R10 media (RPMI, 10% FBS, 1% L-Glu, 1% Pen/Strep) at 4° C in a 1.5 ml eppendorf tube. The needle was rinsed with media 2-3 times to remove remaining cells. The samples were placed on ice or into a cooling rack for transport to the laboratory.

Regarding the donors, there were no major complications following LN biopsy. No subject developed a hematoma or infection. Needle access site bleeding, though expected, was uncommon and resolved with compression. Minor complications were rare, but are known risks of any intervention involving needle placement. One subject developed a mild vasovagal reaction post biopsy that resolved spontaneously after laying flat for 30 minutes and one subject experienced radiating leg pain that resolved with ibuprofen in less than 48hrs.

### LN FNA sample processing

LN FNAs were washed in 15ml cold R10 (1200rpm for 10 minutes). Samples were lysed with 3ml of ACK lysis buffer (Thermo Fisher Scientific A1049201) for 5 minutes at room temperature, then washed with R10, at which point a cell count was taken. If samples were observed to still have significant RBC contamination, those samples were re-lysed with ACK lysis buffer. Cells were washed with R10 media in preparation for flow cytometry Ab staining.

### Criteria for successful LN FNA

Multiple factors should be consider to judge the success of LN FNA sampling including: 1) low granulocyte frequency (>10%), 2) higher total B cell frequency (~20%) than found in blood, 3) presence of LN-specific cell types such as GC-T_FH_ cells and GC B cells, 4) total cell yield (>50,000 cells).

### Blood collection and processing

Standard blood collection by a qualified professional CTRI staff member (MD, nurse, or phlebotomist) at CTRI. The LJI Blood Processing Core processed the blood sample using a standard ficoll separation method.

### Flow cytometry

LN FNA and PBMC samples were transferred to a round bottom 96-well plate and stained with flow a cytometry Ab panel (see below) for 1 hour at 4°C. Cells were then washed twice with cold FACS buffer (washes performed at 2000rpm for 1 minute), fixed with 1% formaldehyde at 4°C for 20 minutes, then washed twice with FACS buffer. Samples were acquired on a BD Fortessa, BD Celesta, or BD Aria II. Flow cytometry data was analyzed using Flow Jo v9 and v10. Anti-human CD8 (RPA-T8, Thermo Fisher) was included in some staining panels.

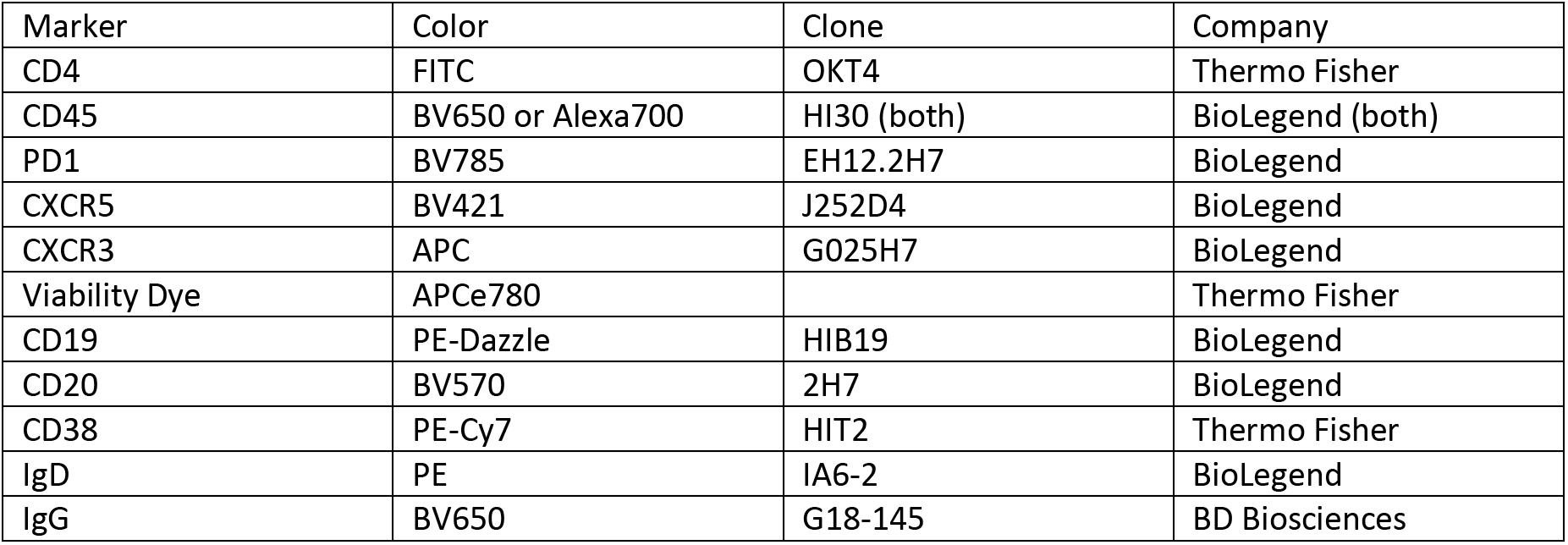
GC cell Panel

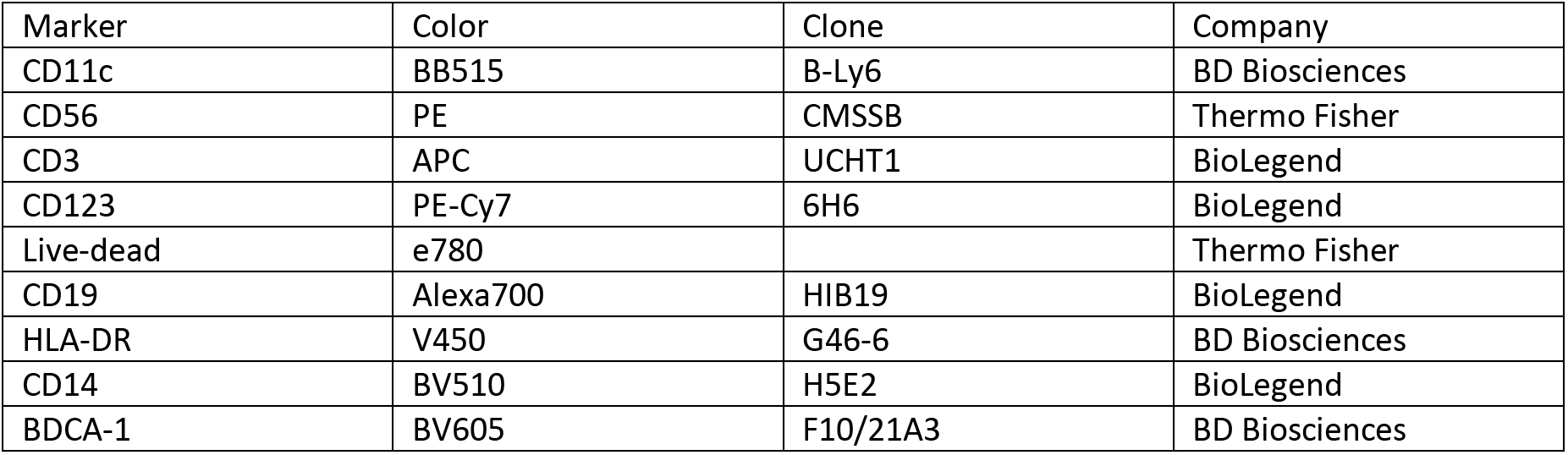
DC and NK cell panel

### Statistical analyses

Prism 7 (GraphPad) was used for statistical analysis. A two-tailed Wilcoxon matched-pairs signed rank test was used for comparison of data in Figure 4D.

### Study approvals

LN FNAs and blood draw protocols were approved by the UCSD Institutional Review Board and accepted by the La Jolla institute Institutional Review Board under a reliance agreement covering the approved protocols. Written informed consent was received from all participants prior to study inclusion.

## AUTHOR CONTRIBUTIONS

C.H.-D. and S.C. designed and supervised the study.

I.N. and S.Y. performed LN FNAs.

S.M.R. and M.S. collected data and performed experiments.

S.M.R., M.S., C.H.-D., and S.C. analyzed data sets.

C.H.-D., S.M.R., I.N., and S.C. wrote the manuscript.

## ACKNOWLEDGEMENTS

We thank the La Jolla Institute (LJI) Clinical Core, especially Gina Levi and Brittany Schwan for expert coordination of the study. We thank Matthew Lariviere and Krystal Caluza of the LJI Blood Processing Core. We also thank the UCSD Center for Translational Research Institute (CTRI) for assistance with LN FNAs and blood draws; the project described was partially supported by the National Institutes of Health, Grant UL1TR001442 of UCSD CTSA. We thank the Flow Cytometry Core at the La Jolla Institute for Immunology for expert cell sorting. This work was supported by the NIH NIAID UM1 Al100663 and UM1 AI144462 to the Scripps CHAVI-ID / CHAVD (SC), and NIH CCHI U19 AI142742. Funding support was also provided by the Human Vaccine Project (SC). The FACSAria II Cell Sorter was acquired through the NIH Shared Instrumentation Grant (SIG) Program (S10 RR027366).

