## Supplemental Information for "Normal human lymph node T follicular helper cells and germinal center B cells accessed via fine needle aspirations"

**Figure S1. Live T and B cells recovered from healthy human LN FNA samples with minimal blood cells**

**Figure S2. NK and dendritic cell populations recovered by LN FNA.**

**Figure S3. Variables affecting LN FNA sampling**

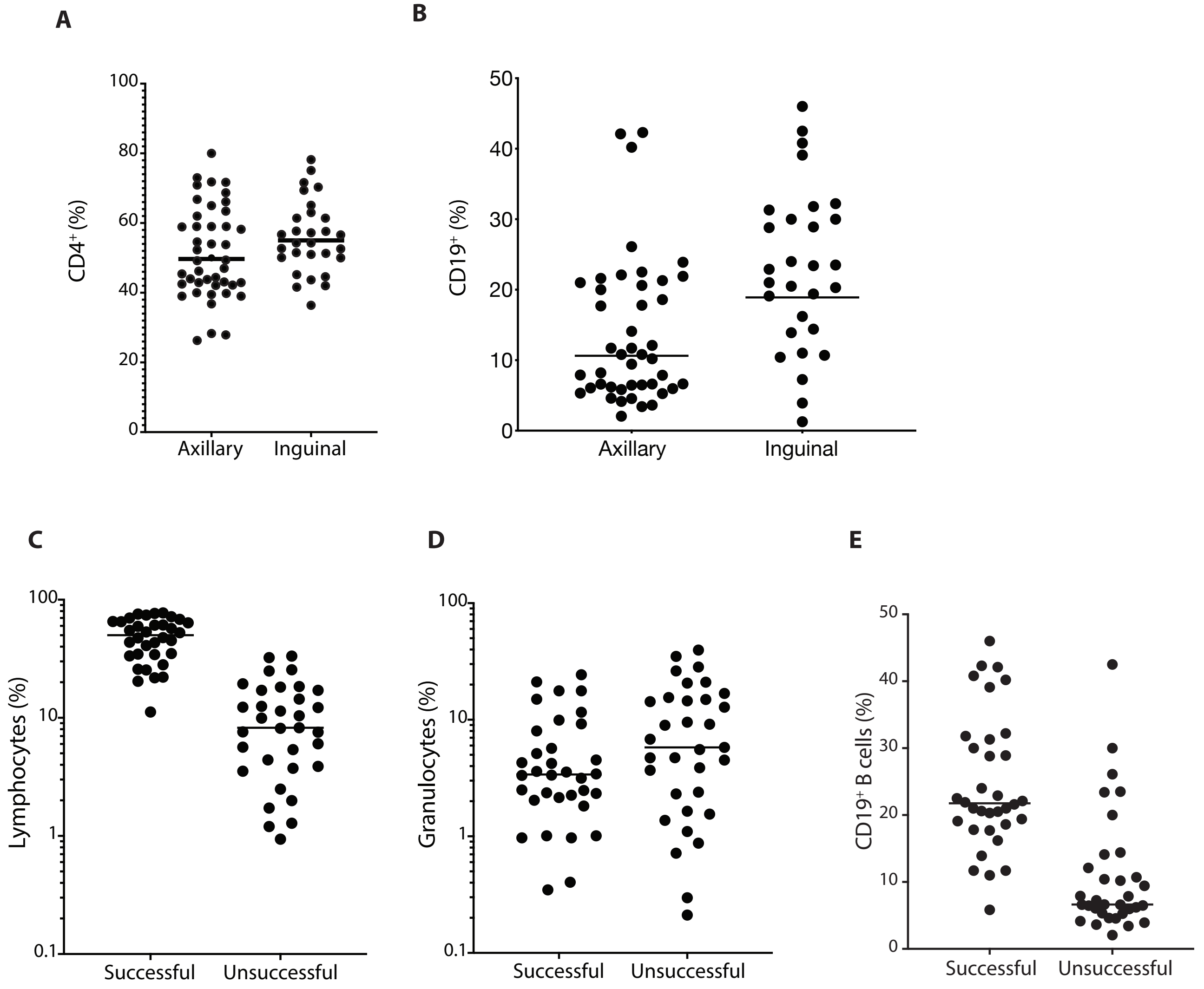


**Figure S1. Live T and B cells recovered from healthy human LN FNA samples with minimal blood cells**

A) Quantification of $CD4^+$ T cell percentages of live lymphocytes. All samples by LN site.

B) Quantification of $CD19^+$ B cell percentages of live lymphocytes. All samples by LN site.

C) Quantification of live CD45+ lymphocytes. All samples by LN FNA success.

D) Graph of granulocyte percentage of total acquired events. All samples by LN FNA success.

E) Quantification of CD19+ B cell percentages of live lymphocytes. All samples by LN FNA success.

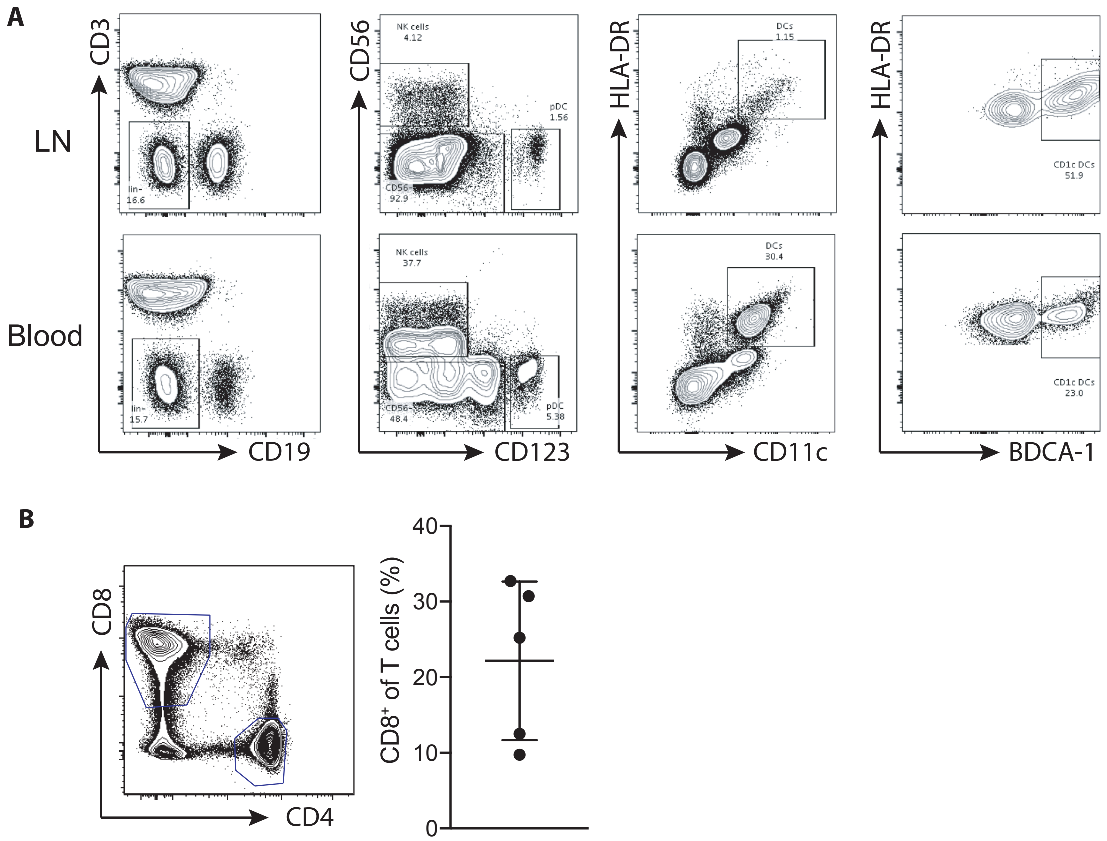


**Figure S2. CD8, NK, and dendritic cell populations recovered by LN FNA.**

A) Flow cytometric identification of human DC and NK subsets in LNs and blood. n=3 LN and n=3 blood samples.
B) Identification of human CD8 T cells in LNs.

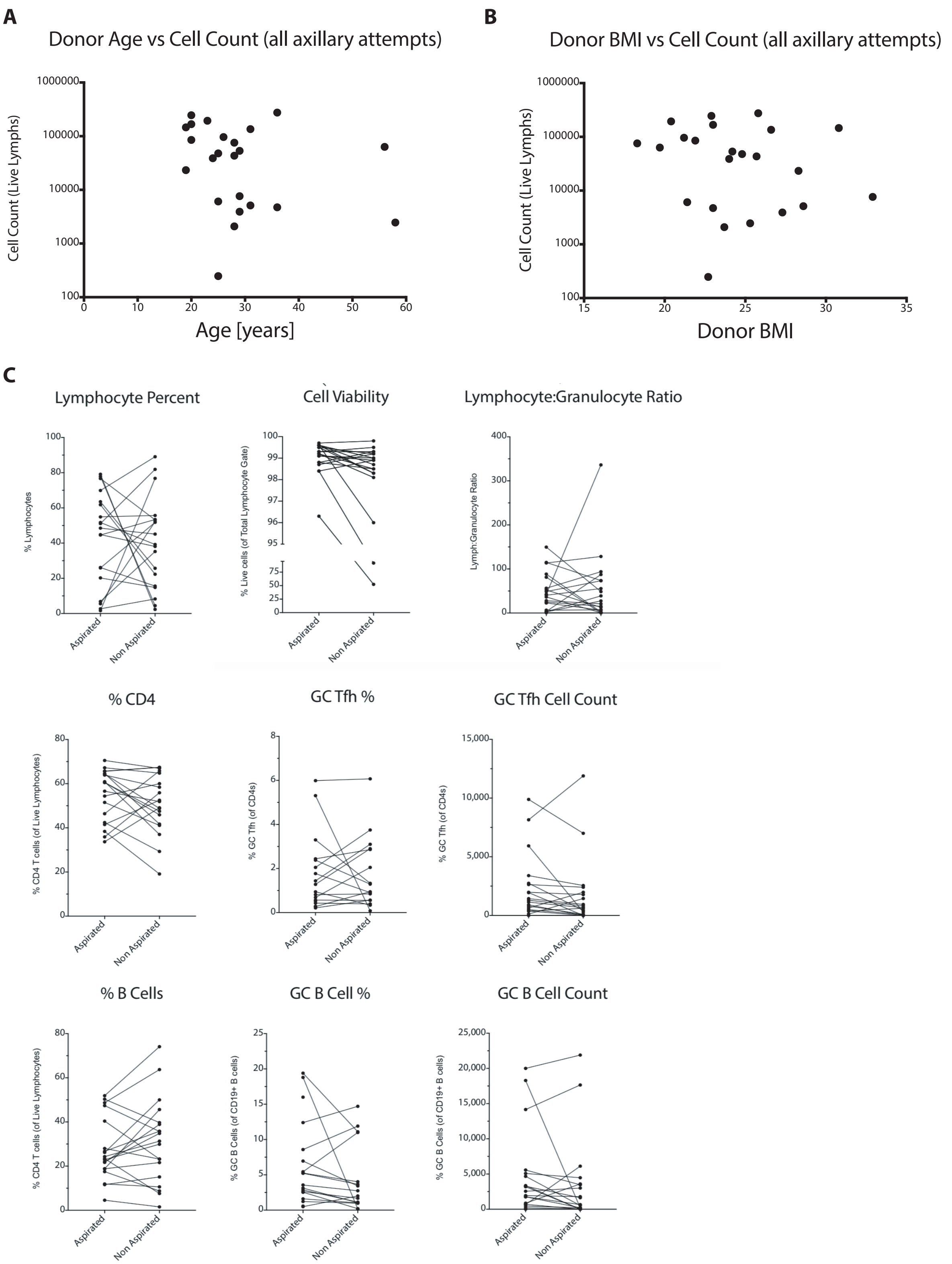


**Figure S3. Variables affecting LN FNA sampling**

A) Cell yield of live lymphocytes versus donor age. All samples.
B) Cell yield of live lymphocytes versus donor BMI. All samples.
C) LN FNA characterization with or without aspiration.
